# IgA-deficient humans exhibit gut microbiota dysbiosis despite production of compensatory IgM

**DOI:** 10.1101/446724

**Authors:** Jason R Catanzaro, Juliet D Strauss, Agata Bielecka, Anthony F Porto, Francis M Lobo, Andrea Urban, Whitman B Schofield, Noah W Palm

**Affiliations:** Section of Pulmonology, Allergy, Immunology, and Sleep Medicine, Department of Pediatrics, Yale School of Medicine, New Haven, CT, USA; Department of Immunobiology, Yale School of Medicine, New Haven, CT, USA; Artizan Biosciences, New Haven, CT, USA; Section of Pediatric Gastroenterology, Department of Pediatrics, Yale School of Medicine, New Haven, CT, USA; Section of Rheumatology, Allergy and Clinical Immunology, Department of Internal Medicine, Yale School of Medicine, New Haven, CT, USA; Section of Pediatric Endocrinology, Department of Pediatrics, Yale School of Medicine, New Haven, CT, USA

## Abstract

Immunoglobulin A is the dominant antibody isotype found in mucosal secretions and enforces host-microbiota symbiosis in mice, yet selective IgA-deficiency (sIgAd) is the most common primary immunodeficiency in humans and is often described as asymptomatic. Here, we determined the effects of IgA deficiency on human gut microbiota composition and evaluated the possibility that secretion of IgM can compensate for a lack of secretory IgA. We used 16S rRNA gene sequencing and bacterial cell sorting to evaluate gut microbiota composition and IgA or IgM coating of the gut microbiota in 15 sIgAd subjects and 15 matched controls. Although sIgAd subjects secreted a significant amount of IgM into the intestinal lumen, this was insufficient to fully compensate for the lack of secretory IgA. Indeed, sIgAd subjects displayed an altered gut microbiota composition as compared to healthy controls, which was characterized by a trend towards decreased overall microbial diversity and significant shifts in the relative abundances of specific microbial taxa. While IgA targets a defined subset of the microbiota via high-level coating, compensatory IgM binds a broader subset of the microbiota in a less targeted manner. We conclude that IgA plays a critical and non-redundant role in controlling gut microbiota composition in humans and that secretory IgA has evolved to maintain a diverse and stable gut microbial community that promotes human health, enhances resistance to infection, and is resilient to perturbation.

## INTRODUCTION

Secretory immunoglobulin A is the dominant antibody isotype found in mucosal secretions and plays an essential role in defense against pathogenic infections.^1^ However, even the absence of infection, the body produces approximately four grams of IgA daily, more than all other antibody isotypes combined.^2^ Much of this IgA is secreted into the intestinal lumen, where it binds to and ‘coats’ specific members of the gut microbiota—the trillions of bacteria that constitutively colonize the human intestinal tract.^3^

Secretory IgA plays a crucial role in shaping the composition, activity and function of the gut microbiota in mice.^4 5^ Various genetically-modified mouse models that display defective secretory antibody responses exhibit dramatic alterations in gut microbial composition and function.^6 7 8 9 10^ However, the role of IgA in shaping the composition and function of the microbiota in humans remains largely unexplored.

Selective IgA deficiency (sIgAd), defined as a serum IgA concentration of less than 7mg/dl with normal levels of serum IgG and IgM in subjects greater than four years of age, is the most common primary immune deficiency in humans. Its incidence varies by geographical region, ranging from 1:700 in those of European descent to 1:18,500 in Japan.^11^ While IgA deficiency is often described as “asymptomatic,” sIgAd subjects exhibit increased incidences of infectious, allergic and autoimmune disorders including inflammatory bowel disease.^12 13^ Nonetheless, the lack of consistent intestinal pathologies in sIgAd subjects suggests that additional immunological defense mechanisms may compensate for the lack of IgA-mediated defenses at mucosal surfaces.

Here, we directly examine the effect of IgA deficiency on the human gut microbiota by defining the composition and stability of the gut microbiota in sIgAd subjects and controls. We confirm previous suggestions that secretory IgM can partially compensate for the lack of secretory IgA in sIgAd patients.^14^ However, compensatory IgM in IgA-deficient subjects displays a distinct microbial binding pattern as compared to conventional secretory IgA and sIgAd subjects exhibit significant alterations in gut microbial community composition. Together, these studies reveal a unique and non-redundant role for secretory IgA in shaping gut microbial community structure in humans.

## RESULTS

### sIgAd subjects exhibit compensatory secretion of microbiota-targeted IgM

Fifteen subjects with reported sIgAd and fifteen healthy controls were recruited by mining electronic medical records and subjected to serum immunoglobulin analyses (Table 1). All subjects with reported IgA deficiency displayed serum IgA levels below the lower limit of detection at the time of study enrollment, but exhibited age appropriate serum IgG titers (Fig 1a). Six of the 15 sIgAD subjects had self-described increased incidences of upper respiratory infections compared to their peers, though none of the IgA deficient subjects had received antibiotics in the 4 weeks prior to study participation. One IgA deficient subject had chronic diarrhea and nonspecific gastrointestinal discomfort without endoscopic evidence of Celiac disease or IBD, and three IgA deficient subjects had a previously diagnosed autoimmune disease (two with type 1 diabetes mellitus and one with Celiac disease).

**Fig 1.**
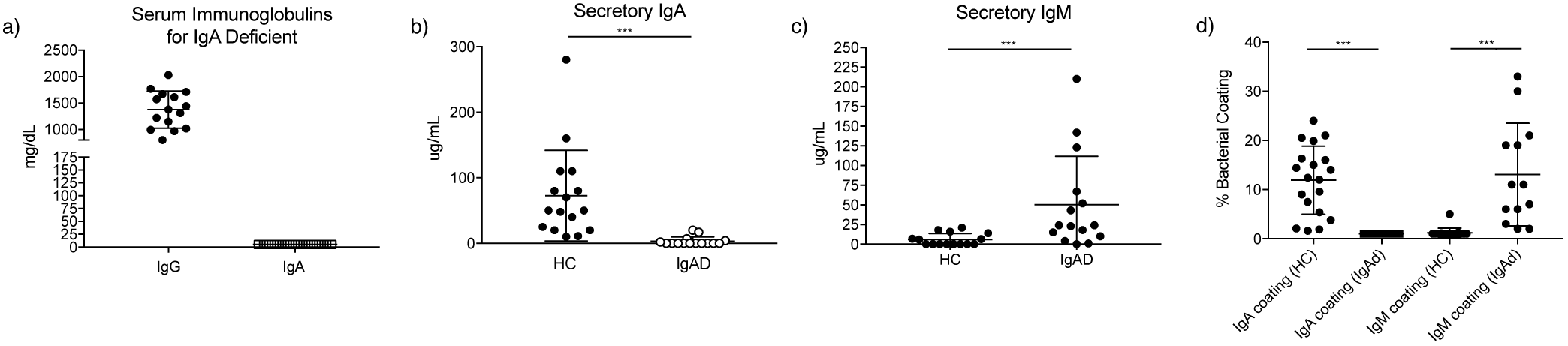
sIgAD subjects exhibit compensatory secretion of microbiota-targeted IgM. (a) Serum IgG and IgA levels for 15 IgA deficient subjects. (b) Secretory IgA levels in the feces as measured by ELISA. (c) Secretory IgM levels in the feces as measured by ELISA. (d) Bacterial coating with IgA or IgM in fecal microbiotas from fifteen healthy controls or sIgAD subjects. ***p<0.001 via Mann-Whitney test

**Table 1.** Baseline characteristics of sIgAD and healthy subjects. Continuous and discrete variables were compared using the Mann-Whitney U test.

|  | IgA deficient | HC | p value |
| --- | --- | --- | --- |
| Total subjects | 15 | 15 |  |
| Mean age | 31.7 | 29.7 | 0.66 |
| Female Sex | 11 (73%) | 10 (67%) | 0.99 |
| Recurrent Infections | 6 (40%) | 0 |  |
| Autoimmune disease | 3 (20%) | 0 |  |
| • T1DM | 2 (13%) | 0 |  |
| • Celiac Disease | 1 (7%) | 0 |  |

sIgAd is normally diagnosed based on the absence of detectable IgA in the serum. Thus, we first examined whether patients with serum IgA deficiency also lacked IgA in their stool. Subjects lacking serum IgA had on average 3.3 ± 1.7 μg/mL of secretory IgA in their stool compared to 72.8 ± 17.8 μg/mL in healthy controls (Fig 1b). The polymeric Ig receptor, which is responsible for transporting IgA across the epithelium into the gut lumen, can also bind to and transport pentameric IgM. Thus, it has been suggested that IgM may at least partially compensate for the absence of IgA in patients with selective IgA deficiency.^14^ In line with this prediction, we found that sIgAd subjects had significantly higher levels of IgM in their stool than controls (mean 50.4 ± 15.8 μg/mL in sIgAD versus 5.9 ± 1.9μg/mL in controls) (Fig 1c). The potential effect of secretory antibodies on the microbiota can also be measured by flow cytometric analysis of IgA or IgM coating of the gut microbiota (Supplemental Fig 1). ^3^ sIgAd subjects displayed undetectable IgA coating by flow cytometry while a significant fraction of bacteria in healthy controls was measurably coated with IgA (1% ± 0.2 versus 11.9% ± 1.8) (Fig 1d). In contrast, sIgAd subjects showed significant bacterial coating with IgM (11.7% ± 2.7) which essentially mirrored the level of IgA coating seen in controls; in contrast, healthy controls exhibited minimal coating with IgM (1.3% ± 0.3) (Fig 1d).

### sIgAd subjects harbor altered gut microbial communities

To assess the effect of IgA deficiency on gut microbial community composition, we performed 16S rRNA gene sequencing on fecal samples from thirteen sIgAd subjects and thirteen controls. IgA has been suggested to support enhanced overall gut microbial diversity in mouse models.^22^ Thus, we first evaluated microbiota diversity in sIgAd subjects versus controls using a variety of metrics. We observed a trend toward decreased diversity in sIgAd subjects for all alpha diversity metrics (observed OTU, Shannon Diversity Index, Shannon and Chao1), but none of these differences reached statistical significance (p<0.05) (Fig 2a). Beta diversity analyses (Principal coordinate analysis based on weighted and unweighted Unifrac distances) revealed that microbial communities in sIgAd subjects occupied an overlapping but distinct cluster as compared to controls (Fig 2b). We next used linear discriminant analysis effect size (LEfSe)^18 19^ to determine whether the relative abundances of specific bacterial taxa differed between sIgAd subjects and controls. At the phylum level, there was no statistically significant difference between the gut microbiota of sIgAd subjects and healthy controls (Fig 2c, Supplemental Fig 2). However, LEfSe analyses at all taxonomic levels revealed five species-level taxa that were significantly more abundant (p<0.01) in sIgAd subjects: *Ruminococcus gnavus* and *bromii*, *Ruminococcus* other, *Eubacterium dolichum*, and an unclassified (UC) *Enterobacteriaceae* (Fig 2c,d). In addition, three taxa were significantly less abundant in sIgAd subjects as compared to healthy controls (p<0.01): UC *Paraprevotella*, *Eubacterium bioforme* and UC *Veillonellaceae* (Fig 2d,e, Supplemental Fig 3). *Prevotella copri* fell just below the stringent significance value of p<0.01, but was notably below the limit of detection in all IgA deficient subjects and was readily detectable in the majority of controls (Fig 2d). Additional differentially abundant taxa were also apparent when we used a less-stringent significance cutoff (p<0.05) (Supplemental Fig 4, 5). Taken together, these results suggest that the intestinal microbiota of subjects lacking secretory IgA is significantly altered as compared to controls.

**Fig 2.**
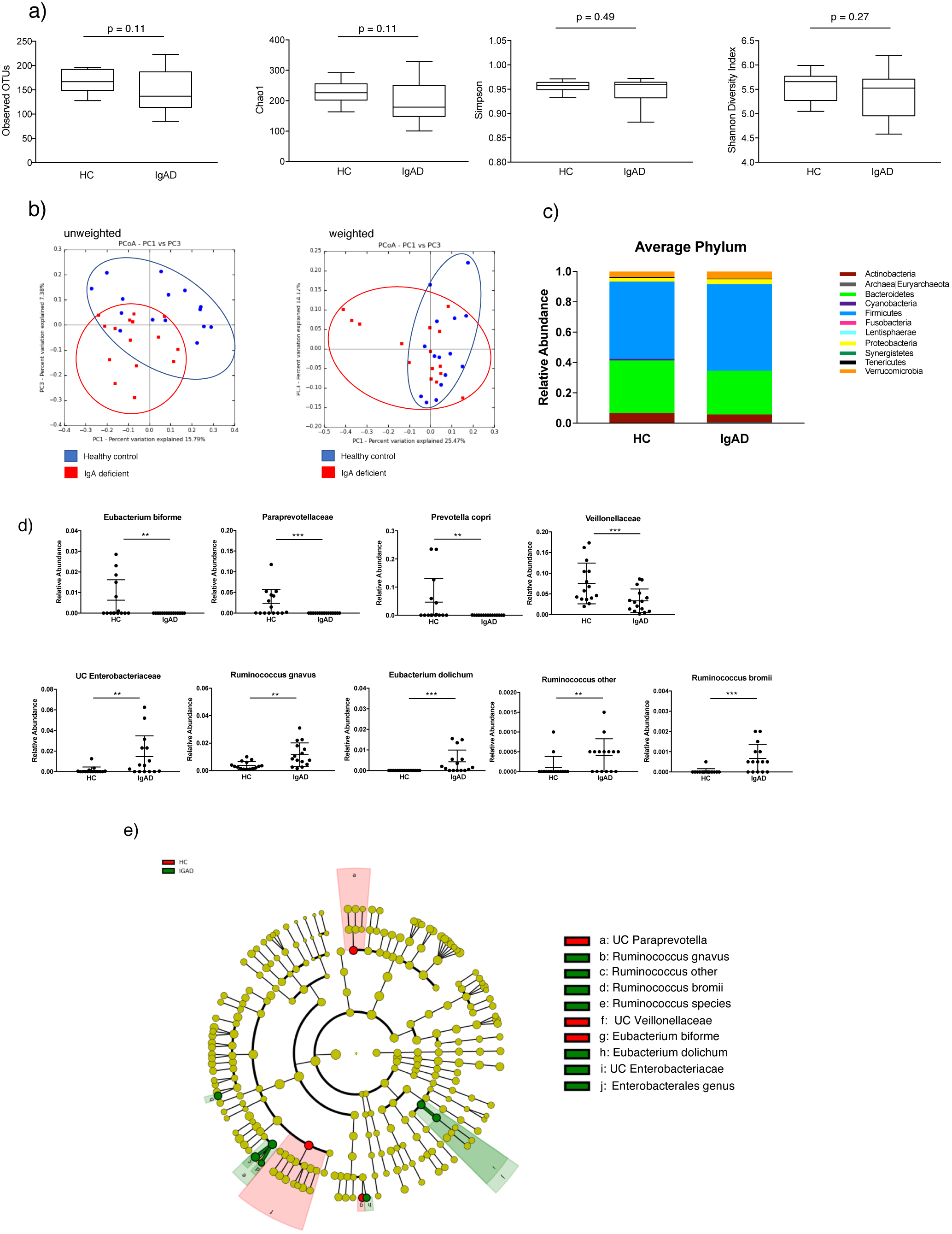
Humans with selective IgA-deficiency harbor altered gut microbial communities. (a) Alpha-diversity analyses in sIgAD subjects and healthy controls (HC) as measured by observed species, Chao1, Simpson and Shannon Diversity indices. (b) Principal Coordinate Analysis plots of unweighted and weighted UniFrac distances. (c and d) Relative abundances of bacterial phyla (c) or species-level taxa exhibiting a significant (p<0.01) difference between sIgAD and control subjects (d). (e) Cladogram displaying linear discriminant analysis effect size (LEfSe) comparisons of sIgAD and HC gut microbial communities. *p<0.01, **p<0.01, ***p<0.001 by Mann-Whitney or LEfSe.

### IgA and compensatory IgM display distinct patterns of bacterial targeting

We previously developed a technology called IgA-Seq^15^ that combines bacterial cell sorting with 16S rRNA gene sequencing to determine the relative levels of IgA coating of the gut microbiota in a taxa-specific manner. We thus performed IgA-Seq or IgM-Seq on fecal bacteria from sIgAd subjects or controls to identify which bacterial taxa were coated with either IgA (in controls) or IgM (in IgA-deficient subjects). To assess the overall distribution of Ig coated versus non-coated taxa, we first evaluated the alpha diversity of the immunoglobulin coated and non-coated fractions in all groups (Fig 3a). We observed that the non-coated fractions displayed a significantly higher species richness (as measured by observed OTUs and Chao) than the antibody-coated fractions in both IgA-deficient subjects and controls. Shannon and Simpson alpha diversity indices, which account for evenness as well as richness, were also significantly lower in the IgA-coated fraction versus non-coated fraction in healthy controls, suggesting that IgA coating in healthy individuals is highly targeted towards a limited subset of gut microbes. In contrast, bacteria coated with compensatory IgM in IgA-deficient subjects showed similar alpha diversity as compared to non-coated bacteria when quantified using the Shannon or Simpson indices. This suggests that compensatory IgM targets a broad spectrum of bacterial taxa, while secretory IgA is more targeted towards specific species or strains. Beta diversity analyses (PCoA of weighted and unweighted Unifrac distances) of all samples performed in triplicate also revealed that IgA‐ versus IgM-coated microbial populations in healthy controls versus IgA deficient subjects clustered separately (Fig 3b). In contrast, non-coated communities in IgA deficient subjects versus controls were largely overlapping (Supplemental Fig 6).

**Fig 3.**
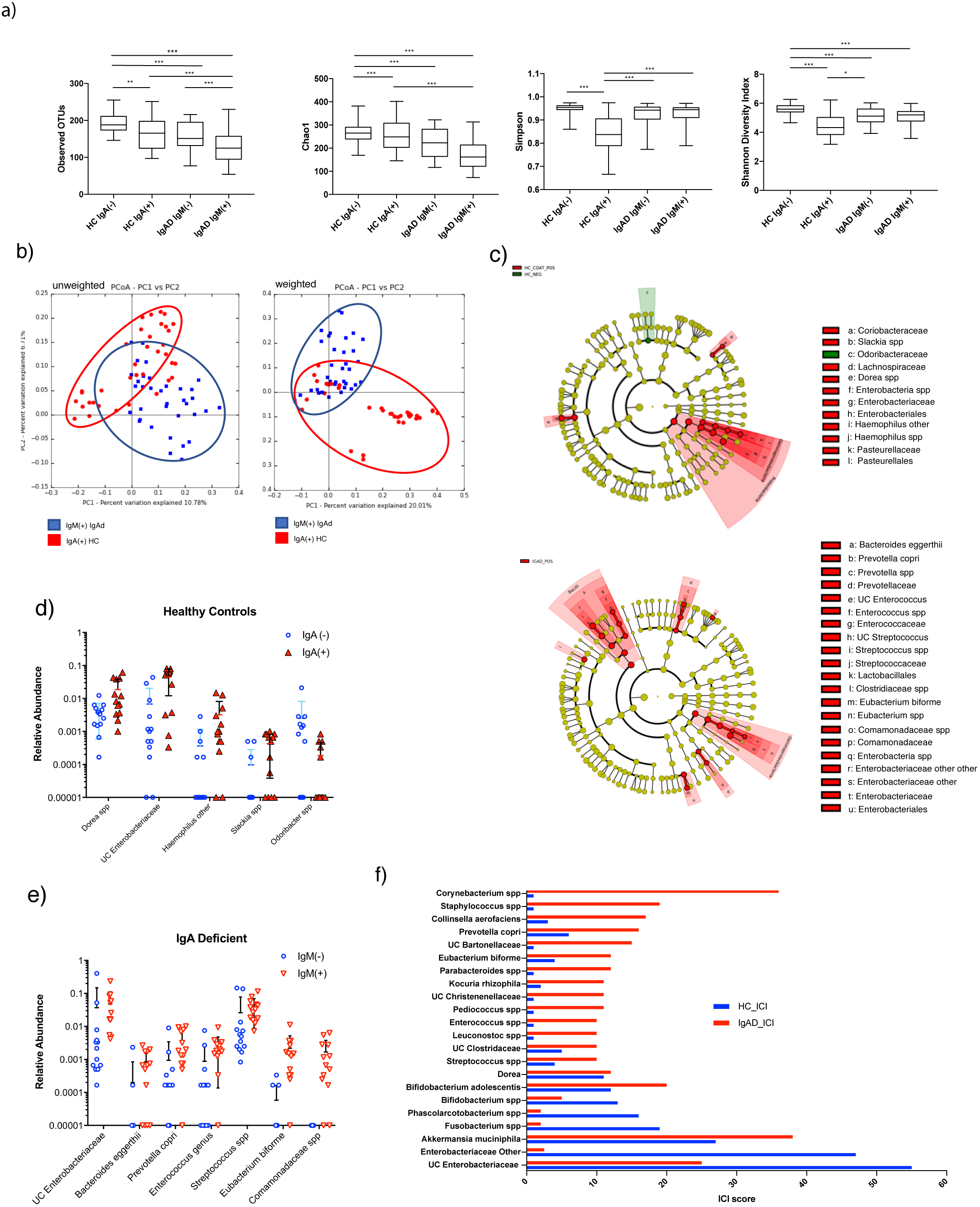
Secretory IgA in healthy controls and compensatory secretory IgM in sIgAD subjects display distinct patterns of bacterial targeting. (a) Alpha diversity analyses (Chao1, Simpson and Shannon Diversity Indices) of antibody coated and non-coated fecal bacteria from healthy controls and sIgAD subjects. (b) Principal Coordinate Analyses of unweighted and weighted UniFrac distances for antibody coated and non-coated bacteria from healthy control subjects and sIgAD subjects. (c) Cladogram of discriminant analysis effect size (LEfSe) comparisons of secretory antibody coated versus non-coated bacteria from healthy controls (coated with IgA or non-coated) or sIgAD subjects (coated with IgM or non-coated). (d) Relative abundances of taxa that were significantly different between IgA coated and uncoated microbes from control subjects (d) or between IgM coated and uncoated sIgAD subjects. (e) Bar chart depicting IgA or IgM coating index scores (ICI Scores) of taxa with ICI scores > 10 in either sIgAD subjects or healthy controls.

To identify which specific taxa were targeted by IgA or IgM, we next compared the taxonomic compositions of the Ig-coated versus non-coated fractions in IgA-deficient subjects or controls using LEfSe (Fig 3d,e). These analyses revealed that *Haemophilus other*, *Dorea spp*, *Slackia spp* and *UC Enterobacteriaceae* were significantly IgA coated in healthy controls (enriched in the IgA coated fraction versus non-coated fraction; p<0.01), while only *Odoribacter spp* was significantly enriched in the non-coated fraction (Fig 3c,d). In contrast, multiple taxa were significantly coated with IgM (p<0.01) in IgA deficient subjects, including *Prevotella copri*, *Bacteroides eggerthii*, an *Enterococcus genus*, *Streptococcus spp*, *Eubacterium bioforme*, *Comamonadaceae spp* and an *UC Enterobacteriaceae* (Fig 3c,d). Only UC *Enterobacteriaceae* was significantly coated (p<0.01) in both controls and IgA deficient subjects (Fig 3e). To quantify and directly compare the relative levels of secretory antibody coating between cohorts, we calculated an IgA or IgM coating index (ICI Score) for both populations by dividing the relative abundance of a given taxon in the coated fraction by the relative abundance in the non-coated fraction (ICI = relative abundance (IgA+ or IgM+)/relative abundance (IgA‐ or IgM‐)). We found that eighteen taxa in the IgA deficient cohort had ICI scores greater than or equal to 10, while only eight taxa in the controls had similarly high ICI scores (Fig 3f).

### sIgAd subjects display normal gut microbiota stability

Ecological models of community stability and resilience suggest that increased diversity is associated with increased overall stability and studies in mice suggest that secretory IgA can promote a more diverse microbial community.^2 3^ To begin to characterize the stability of the fecal microbiota in sIgAd subjects, we collected repeat fecal samples from sIgAd subjects and controls 6-10 months after our original sample collection. Eight sIgAd subjects and seven control subjects provided repeat samples. All subjects and controls reported no exposure to antibiotics one month prior to donation of the follow up sample. Comparison of the taxonomic compositions of original and repeat samples by LEfSe revealed comparable microbiota stability in both sIgAd subjects and controls. Indeed, using a p value cutoff of 0.01, there were no significant differences between original and repeat samples in either group; using a less stringent p value cutoff of p<0.05, only two taxa, *Coprococcus* spp and *Dorea spp*, were more abundant in the initial sample while a single taxon, *UC Bacteroides*, was more abundant in the second sample (Fig 4a). Notably, *Prevotella copri*, *Paraprevotella spp* and *Eubacterium bioforme* remained below the limit of detection in sIgAd subjects upon repeat sampling, but were still present in the majority of controls. This suggests that alterations in the relative abundance of these taxa in IgA-deficient subjects versus controls are durable and consistent over time. Finally, we compared initial and repeat sampling of microbial communities from sIgAd subjects and controls using a variety of alpha diversity metrics (Fig 4b). These analyses demonstrated that alpha diversity was largely consistent between initial and repeat microbiota collections for both IgA deficient subjects and controls. However, sIgAd subjects exhibited significantly lower alpha diversity than controls (p<0.05) for both the initial and repeat samples as measured via Shannon and Simpson indices. Finally, we measured intra-individual microbiota stability over time using the Bray-Curtis (BC) dissimilarity metric. Overall, we observed that sIgAd subjects and controls displayed similar mean BC dissimilarity over time, though IgA deficient subjects demonstrated greater intra-individual variability in dissimilarity between initial and repeat samples as compared to healthy controls (Fig 4c).

**Fig 4.**
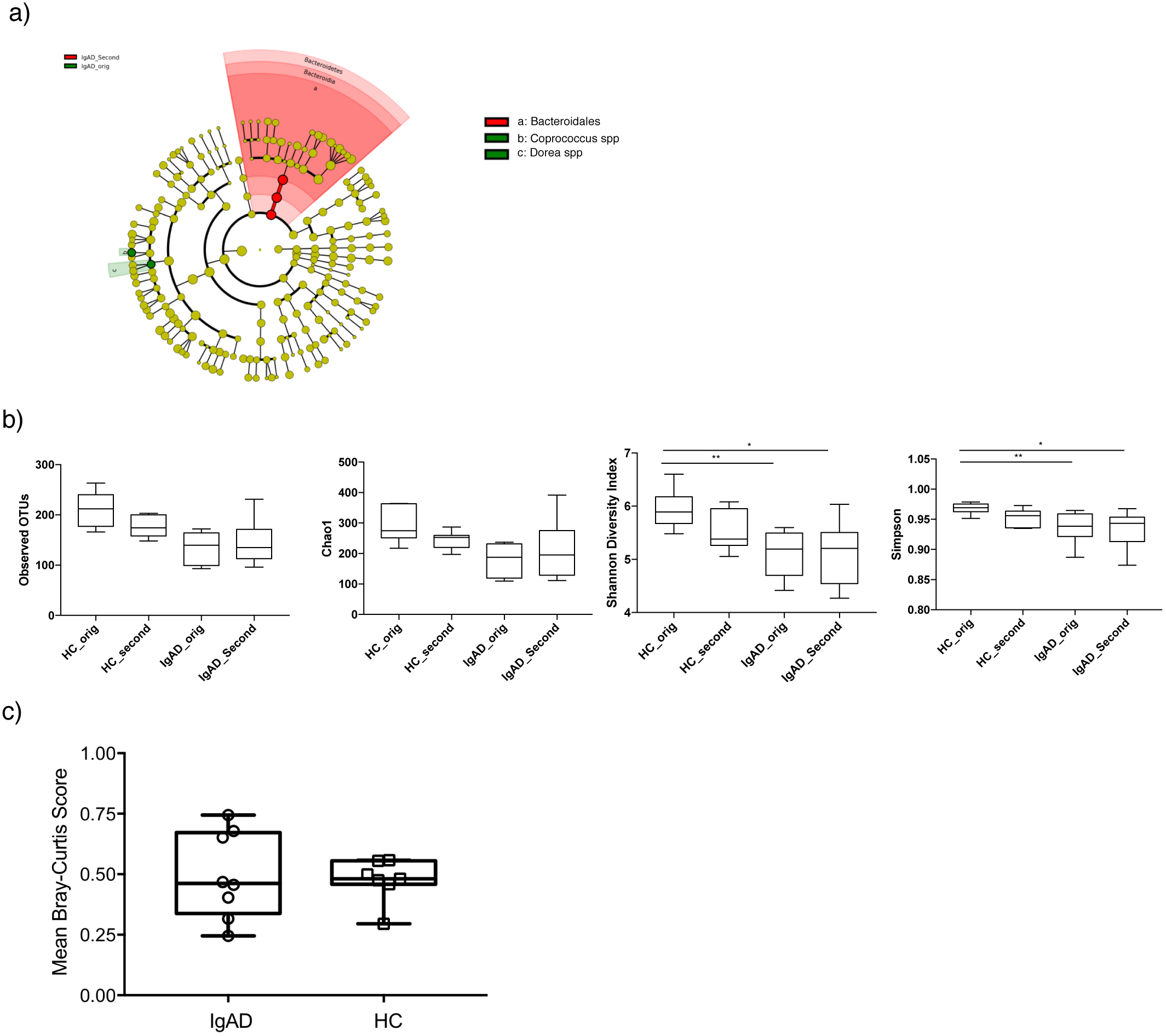
IgA-deficient subjects display normal gut microbiota stability. (a) Cladogram generated from linear discriminant effect size (LEfSe) analyses comparing initial and repeat samples from sIgAD subjects. (b) Alpha diversity analyses of gut microbiota composition comparing initial and follow up stool samples from both healthy controls and sIgAD subjects. (c) Bray-Curtis comparisons of initial and repeat samples from healthy control and sIgAD subjects.

## DISCUSSION

Immunoglobulin A is best known for its role in protecting against infections at mucosal surfaces; however, it has recently become clear that IgA also plays a critical role in shaping the composition and function of the gut microbiota in mice. Selective IgA deficiency is the most common primary human immunodeficiency, which raises the question of whether IgA also plays a critical role in shaping gut microbial communities in humans. We addressed this question by examining gut microbiota composition and bacterial targeting by IgA and IgM in sIgAd subjects versus controls.

As IgA deficiency is largely asymptomatic, many patients with sIgAd go undiagnosed. To obtain a collection of subjects that was broadly representative of the IgA-deficient community at large, we used the electronic medical record system at Yale to identify IgA deficient patients from the broader New England population. These patients’ IgA deficiencies were uncovered by means typical for this population—*e.g.,* evaluation for recurrent infections or gastrointestinal disorders, or screening for Celiac disease. Thus, our recruitment strategy captured a population of sIgAd subjects that is representative of the sIgAd population seen in a variety of primary and subspecialty clinics.

We found that the gut microbiota in sIgAd subjects was significantly altered as compared to healthy controls, with a trend towards an overall reduction in alpha diversity as well as defined changes in the presence and abundance of specific taxa. We also found that secretory IgM can at least partially compensate for the lack of IgA since IgM coated the gut microbiota of our IgA-deficient cohort at a level that was, on average, almost identical to the level of bacterial coating with IgA in controls. However, compensatory IgM coating differed from IgA coating in a few important ways. In particular, we found that compensatory IgM targeted a broader population of microbes as compared to IgA, which suggests that compensatory IgM may show less specificity towards specific microbes or epitopes as compared to IgA. These data imply that the secretory IgA response may be, on average, more T cell-dependent and antigen-specific than compensatory IgM responses; notably, IgA-deficiency can result from defective T-B interactions in at least a subset of IgA-deficient subjects.^23^ Our preliminary results also suggest that the longitudinal stability of the microbiota in IgA deficient subjects, while similar on average to our controls, displays greater intra-individual variability. This highlights the possibility that select sIgAd subjects may experience deviations from a healthy microbial composition; such perturbations may play a role in the development or exacerbation of autoimmune or infectious diseases.

IgA is classically thought of as a restrictive factor because of its anti-pathogen effects.^24^ Indeed, we noted that specific taxa that are normally targeted by IgA, such as members of the *Enterobacteriaceae*, were more abundant in the microbiota of sIgAd subjects as compared to healthy controls. However, recent studies have revealed that IgA also can enhance colonization by specific bacterial taxa either indirectly, through restriction of competitors, or by directly enhancing colonization of host-associated niches such as colonic crypts.^22 25^ In line with these findings, we identified multiple taxa that were uniquely present in controls and nearly completely absent (or fell below the limit of detection) in IgA-deficient subjects. We also noted that even related species sometimes displayed apparently divergent effects of IgA coating: *Eubacterium biforme* was observed exclusively in controls and was absent in IgA-deficient subjects, while *Eubacterium dolichum* was only seen in IgA-deficient subjects and not in controls. These data support the idea that IgA can have opposing effects on distinct members of the microbiota. Future studies will be necessary to understand what determines when IgA restricts versus promotes bacterial colonization or growth.

We used IgA‐ and IgM-Seq to determine taxa-specific patterns of targeting of the microbiota by IgA and compensatory IgM. We found that certain taxa, such as *UC Enterobacteriaceae* and *Akkermansia muciniphilia* were highly coated with both IgA and compensatory IgM. In contrast, select taxa were uniquely highly coated with IgA and showed lower levels of coating with compensatory IgM. This raises the possibility that distinct subsets of commensal bacteria may engage with the host in unique ways that lead to these divergent outcomes. The bacterial features or behaviors that define these distinctions will be an important subject of future research.

While we were preparing this study for publication, a report was published detailing an examination of the microbiota of IgA-deficient patients in a European cohort. Similar to our study, Fadlallah *et al.*^26^ also found a role for IgA in shaping gut microbiota composition in humans. However, our data differ in some important ways. First, while Fadlallah et al.^26^ found no evidence for an overall change in alpha diversity between IgA deficient subjects and controls, we observed a strong trend towards decreased alpha diversity when taking into consideration both richness and evenness of the microbial population in IgA deficient patients. Notably, Jorgensen et al.^27^ also recently reported that CVID patients with low levels of serum IgA exhibited reduced gut microbiota alpha diversity.^28^ The specific alterations in the gut microbiota in sIgAd subjects and the patterns of IgA and compensatory IgM coating that we observed also sometimes diverged from the patterns observed by Fadlallah et al.^26^; for example we observed seemingly opposite effects of IgA deficiency on the relative abundance of specific taxa (e.g., *Prevotella copri* and *Veillonellaceae*) as compared to Fadlallah et al.^26^ There are multiple potential explanations for these differences. For example, differences in the clinical history of the subjects could explain some differences: 52% of the IgA deficient individuals in the Fadlallah et al.^26^ cohort presented with autoimmunity, whereas only 20% of our subjects had a known autoimmune condition. In addition, differences in European versus American microbial communities or in the methodologies used to evaluate microbial community composition may also explain some of these differences. Indeed, Fadlallah et al.^26^ used a combination of metagenomics and 16S rRNA gene sequencing for their microbiota analyses, while we used exclusively 16S rRNA gene sequencing. Finally, because of the asymptomatic nature of IgA-deficiency, identifying patients is often challenging and, thus, both our study and Fadlallah et al. evaluated similarly small cohorts of patients. Thus, it is theoretically possible that the differences between these two studies are due to random chance. Future studies employing larger cohorts that include patients from multiple continents will be necessary to definitively identify universal features of microbial dysbiosis in humans with IgA-deficiency.

In conclusion, these studies demonstrate that IgA plays a critical and non-redundant role in shaping the composition of the gut microbiota in humans and IgM can partially, but not completely, compensate for the absence of IgA. Thus, while the relatively high rate of asymptomatic sIgAd has been used to argue against the importance of IgA, our studies highlight the critical role of this secretory antibody in the maintenance of host-microbiota symbiosis.

## METHODS

### Sample collection

Human study protocols were approved by the Institutional Review Board (HIC # 1607018104) of the Yale School of Medicine, New Haven, CT. IgA deficient subjects were identified via the EPIC electronic medical record system and all subjects resided in the state of Connecticut. Demographics, medical history and other clinical variables were collected following enrollment. Healthy subjects were recruited via an advertisement on the Yale campus. Fecal samples were collected at home and stored at −20 °C before overnight shipment to the Palm Lab on ice in an insulated container. Samples were then stored at −80 °C until use. Informed consent was obtained from all subjects.

### Measurement of serum immunoglobulins

Whole blood was drawn by venipuncture into tubes without anticoagulant (BD, USA). Blood was allowed to clot for 30 minutes, centrifuged at 10,000 x g for 10 minutes, and serum supernatant was collected. Serum was probed for IgA and IgM via ELISA (coating Sigma SAB3701393, Detection of IgA1 Southern Biotech 9130-08, IgA2 Southern Biotech 9140-08, IgM Southern Biotech 9020-08).

### Bacterial flow cytometry, IgA-Seq and IgM-Seq analyses

Fecal flow cytometry and IgA-Seq were performed essentially as previously described except that Ig-coated bacteria were isolated using MACS beads and a custom-built 96 well magnet followed by negative selection using MACS multi 96 columns.^15^ Bacteria were stained with either PE-conjugated Anti-Human IgA (1:10; Miltenyi Biotec clone IS11-8E10) or FITC‐ or PE‐ conjugated Anti-Human IgM (1:50 JI 109-096-008). IgM-Seq samples were processed identically to IgA-Seq except that PE-anti-IgM was used in place of PE-anti-IgA.

### 16S ribosomal RNA gene sequencing and microbial community composition analysis

The V4 region of the 16S ribosomal RNA gene was sequenced on a MiSeq (2×250; Illumina) as described previously.^15 16 17^ Microbial diversity analyses were performed using the Quantitative Insights Into Microbial Ecology (QIIME version 1.9) analysis suite. Reads were assembled, demultiplexed and quality filtered with a Q-score cutoff of 30. Open-reference OTU picking against the Greengenes reference database was used to cluster into Operational Taxonomic Units (OTUs) at 97% identity. The Ribosomal Database Project classifier (RDP) and the May 2016 Greengenes taxonomy were used to assign taxonomy to representative OTUs.^18 19 20^ OTUs of less than 0.01% relative abundance, and contaminating OTUs found in water-only controls were removed from OTU tables. QIIME was used for all microbial ecology analyses^18^ and Linear Discriminant Analysis Effect Size (LEfSe)^21^ was used to compare taxonomic abundance between groups.

Contributors. JRC designed and performed experiments, analyzed data and wrote the paper; NWP conceived ideas, wrote the paper and oversaw the research; AB and JDS performed experiments; FML, AFP, AU and WBS provided advice and/or assisted with patient recruitment.

Declarations of interests: WBS and JDS are employees of Artizan Biosciences and are visiting scientists in Immunobiology at Yale. NWP is a co-founder of, consultant for and receives research funding from Artizan Biosciences. All other authors declare no competing interests.

Funding: The funding for these studies was provided by internal Yale sources.

## LEGENDS

Supplemental Fig 1 | Representative flow cytometric analysis of IgA coated bacteria in a healthy subject and IgM coated bacteria in an IgA deficient subject,

Supplemental Fig 2 | Barplots depicting relative abundances of phyla from fifteen sIgAD subjects and healthy controls.

Supplemental Fig 3 | Linear discriminant analysis (LDA) scores for differentially abundant taxa enriched in healthy controls versus sIgAD subjects with p<0.01.

Supplemental Fig 4 | Cladogram generated from linear discriminant effect size (LEfSe) comparisons between healthy controls and sIgAD subjects (p<0.05).

Supplemental Fig 5 | Linear discriminant analysis (LDA) scores for differentially abundant taxa enriched in healthy controls versus sIgAD subjects (p<0.05).

Supplemental Fig 6 | Principal Coordinate analyses of unweighted and weighted UniFrac distances for uncoated taxa from healthy controls and sIgAD subjects.

